# Why are rare variants hard to impute? Coalescent models reveal theoretical limits in existing algorithms

**DOI:** 10.1101/2020.08.10.245043

**Authors:** Yichen Si, Brett Vanderwerff, Sebastian Zöllner

## Abstract

Genotype imputation is an indispensable step in human genetic studies. Large reference panels with deeply sequenced genomes now allow interrogating variants with minor allele frequency < 1% without sequencing. While it is critical to consider limits of this approach, imputation methods for rare variants have only done so empirically; the theoretical basis of their imputation accuracy has not been explored. To provide theoretical consideration of imputation accuracy under the current imputation framework, we develop a coalescent model of imputing rare variants, leveraging the joint genealogy of the sample to be imputed and reference individuals. We show that broadly used imputation algorithms includes model misspecifications about this joint genealogy that limit the ability to correctly impute rare variants. We develop closed-form solutions for the probability distribution of this joint genealogy and quantify the inevitable error rate resulting from the model misspecification across a range of allele frequencies and reference sample sizes. We show that the probability of a falsely imputed minor allele decreases with reference sample size, but the proportion of falsely imputed minor alleles mostly depends on the allele count in the reference sample. We summarize the impact of this error on genotype imputation on association tests by calculating the *r*^2^ between imputed and true genotype and show that even when modeling other sources of error, the impact of the model misspecification have a significant impact on the *r*^2^ of rare variants. To evaluate these predictions in practice, we compare the imputation of the same dataset across imputation panels of different sizes. While this empirical imputation accuracy is substantially lower than our theoretical prediction, modeling misspecification seems to further decrease imputation accuracy for variants with low allele counts in the reference. These results provide a framework for developing new imputation algorithms and for interpreting rare variant association analyses.

## 1 Introduction

Emerging results from sequencing studies elucidate the impact of rare variants on the etiology of complex diseases [1]. Genome sequencing studies with deep coverage allow directly assessing these variants [2], but such studies are still expensive. As an alternative, array-based technologies can be employed at a substantially lower cost. Commercial genotyping arrays cover a pre-selected set of common variants, and in some cases low frequency variants known to be of interest from previous studies. To recover high resolution genetic information, genotype imputation compares assayed genotypes to a sequenced reference panel, thus leveraging the shared genealogy between genotyped (target) individuals and reference panel to infer unobserved genotypes [3].

Modern imputation methods combined with large reference panels have achieved high accuracy among even low frequency variants. For example, using the TOPMed data as reference, the average imputation quality (*r*^2^) of variants with frequency 0.1% is over 0.90 for both African and European ancestry genomes [4]. Such high resolution improves the power of genome-wide association studies (GWAS) for low frequency and rare variants, and enables joint analysis across studies with different sets of genotyped variants [5]: Recently, the TOPMed consortium identified a new risk variant for breast cancer by imputing rare variants with minor allele frequency (MAF) < 0.5% into the UK Biobank [4]. The advent of affordable whole genome sequencing generating large collections of reference haplotypes combined with efficient imputation algorithms will power more of such discoveries.

Most of these modern imputation methods are based on the Li and Stephens’ model [6]. They leverage that haplotypes from unrelated individuals sharing chromosome segments from a common ancestor. These segments are more similar to each other if their common ancestor is more recent. Among haplotypes in a large reference panel, the haplotype that has the most recent common ancestor (MRCA) with the target haplotype probably also tends to have similar genotypes as the target. Using a Hidden Markov Model (HMM), the Li and Stephens’ algorithm models each target haplotype in the study as an imperfect mosaic of haplotypes from the reference panel. The haplotypes making up this mosaic are inferred to be the most closely related to the target, and thus provide information for unobserved genotype information. The HMM framework is computationally tractable and naturally approximates recombination and mutation in its transition and emission probabilities.

Simulation studies have shown that imputation quality depends on the imputed variants’ allele frequency, genomic context, the size of the reference panel and population demographics [7, 5]. Such studies illustrate that, despite significant improvements, imputation of rare variants still has high uncertainty: The average squared correlation (*r*^2^) for variants with MAF 0.01% is below 0.5 even with the largest reference from the same continental population [4]. But simulation studies only provide a limited opportunity to understand these limits. Although they have the flexibility to mimic a particular imputation setting, they are often computationally expensive and hard to generalize. Alternatively, probabilistic models capturing the basic properties of the imputation process can make predictions before data collection and are easily generalizable.

The key in such a probabilistic model is the relatedness between reference and target haplotypes, which explains how the reference informs unobserved genotypes in target haplotypes. Kingman’s coalescent [8] provides a suitable theoretical framework for modelling the shared genealogy of these haplotypes. The coalescent traces the genealogy back in time, modeling a sequence of events where individuals find their common ancestors. Mutations resulting in polymorphisms can be mapped to branches on the coalescence tree at each locus, with all its descendants carrying the derived allele (Figure 1). The coalescent time (the time it takes for two or more haplotypes to find their MRCA, TMRCA) thus gives a measure of expected genetic dissimilarity, since only mutations occurring more recent than the coalescent time can result in different alleles among those haplotypes.

**Figure 1:**
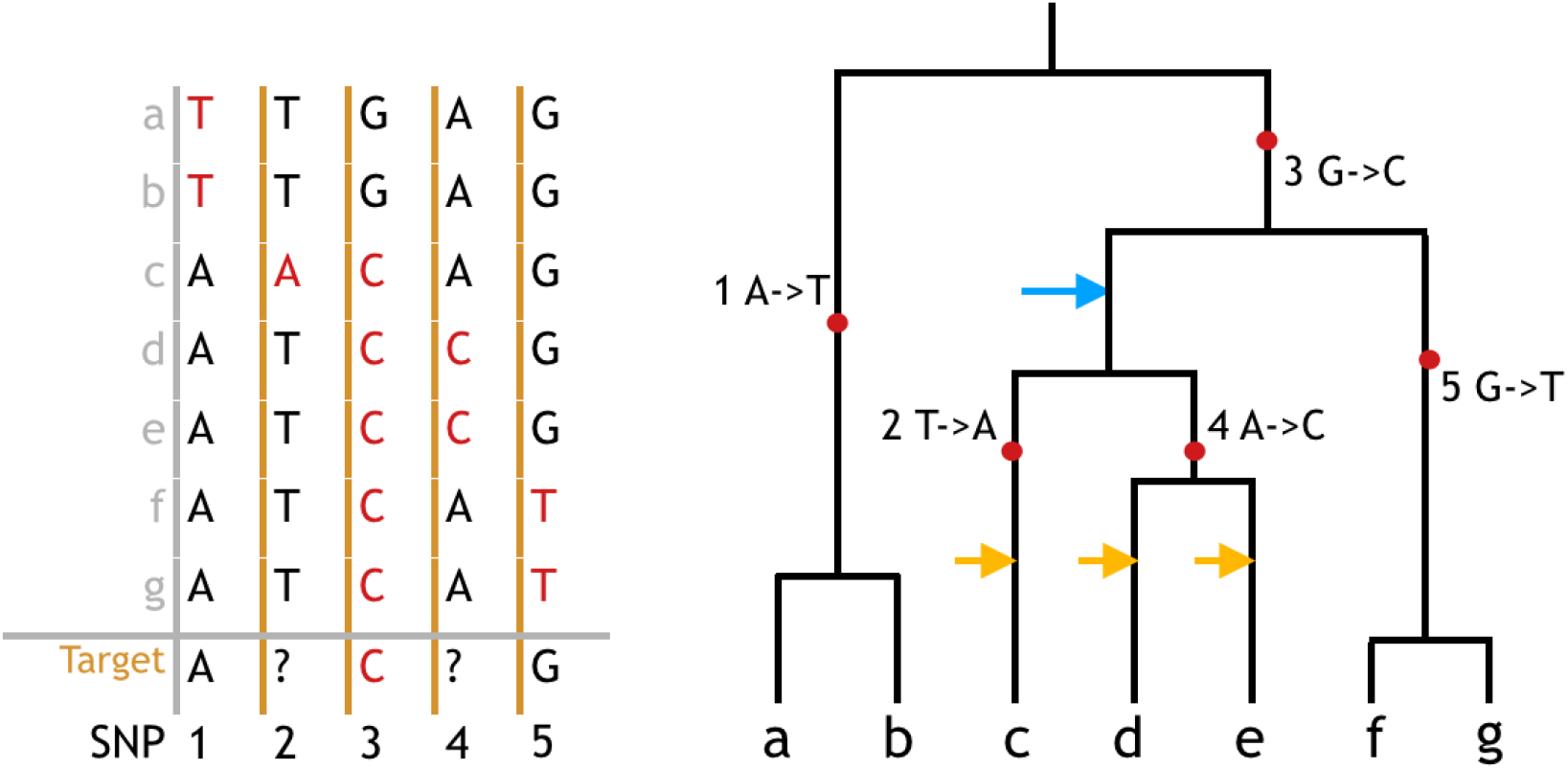
Connection between genotype sequences and coalescence tree. On the left is the genotype matrix of 8 haplotypes (*a* ~ *g* as reference plus a target haplotype) at 5 SNPs (1 ~ 5). Ancestral alleles are colored black, derived (mutated) alleles are colored red. Question marks denote sites where the target is not genotyped. The Li and Stephens algorithm models the the target haplotype as the observation from a HMM where reference haplotypes *a* ~ *g* constitute the state space. By identifying which of the reference haplotypes is the hidden state of the target haplotype, the algorithm can identify which allele to infer for the unknown sites. Note that haplotypes *c, d* and *e* all have the same alleles as the target at the genotyped loci, making them equally likely to be the hidden state without further information. On the right is the corresponding coalescence tree assuming no recombination. Mutations are labeled on the branch, with all the descendants (leaves) carrying the derived allele. Template haplotypes *c, d* or *e* being the hidden state is equivalent to the target haplotype coalescing at one of the yellow arrows. As the Li and Stephens algorithm assumes a single best template, it does not consider the possibility that the target may fist coalesce with the branch pointed by the blue arrow. All indicated coalescence events are compatible with the observed genotypes but give different imputation results.

Some aspects of imputation accuracy have already been explored using coalescent theory. Jewett et al. studied the scenario where the target and reference are sampled from two populations diverged in the past, and derived expected imputation error rate as a function of reference size and divergence time [9]. Huang et al. further included mutation rate and marker density as factors; and analyzed the potential gain in accuracy by choosing the reference panel from a more closely related population[7]. Here we develop a coalescent approach to understand imputation accuracy within population, focused on rare variants. It is useful to recognize that imputation error has two types of sources: (1) failure to identify haplotypes most closely related to the target as the template and (2) true differences between the template and the target haplotype due to recent mutation events. While improving imputation algorithms may reduce error of the first type, error of the second type is a result of the underlying genealogical relation between the target and the reference sample and how this relationship is modelled. Here we focus on modelling the second type of error, where we determine the error that is immanent to the Li and Stephens’ model.

For this purpose, we consider one target haplotype with missing genotype and *n* fully sequenced haplotypes as references. These *n* + 1 observed lineages form the leaves of a binary tree with their (unobserved) ancestors as internal nodes. Intuitively, if the target first finds its MRCA with a set of reference haplotypes, their genotypes are the most similar to the target thus the most informative for imputation. The Li and Stephens’ algorithm assumes that there is exactly one such most closely related reference haplotype, but we show that this assumption is wrong with probability 1/3.

We compute how often this misspecification leads to an ambiguous or wrongly imputed genotype, assuming the imputation algorithm correctly identifies exactly those haplotypes that are most closely related to the target sample in the reference. We then provide the probability of generating a particular imputed dosage conditional on allele frequency of the variant and the size of reference panel. We also quantify the imputation accuracy in terms of the *r*^2^ between the imputed dosage and the true genotype, and show that, as a result of this misspecification, the *r*^2^ largely depends on the allele count in the reference panel, improving only marginally with increased reference size. We assess the impact of population history on these results and use coalescent simulation to confirm our analytic results. Taken together, our approach provides the minimum size of the reference panel necessary to achieve the desired imputation accuracy for a given allele frequency.

We evaluate the upper bounds predicted with this model where the only source of error is model misspecification by imputing a sample of 56,984 individuals using reference panels of different sizes and evaluating the empirical imputation accuracy > 60, 000 variants. We observe substantially higher imputation error in the empirical results than the theoretical bound in our model. Moreover, when conditioning on minor allele count (MAC), imputation error is larger in larger reference panels. We also observe that for MAC 2 ~ 10, imputation error increase much faster with decreasing MAC, consistent with our result that especially for these MACs, model misspecification reduces imputation accuracy.

The model we develop here can also be leveraged to improve current imputation algorithms. For example, most imputation algorithms assume that the distance between switching template haplotypes reflects recombination events and is exponentially distributed [10]. We derive the length distribution of contiguous segments without observable recombination breakpoints, and show that it differs substantially from the exponential assumption by having thicker tails for both very short and long segments.

## 2 Methods

We consider imputing a single target variant for one target haplotype with a reference panel consisting of *n* reference haplotypes. For simplicity, we only discuss biallelic sites denoting the ancestral allele as 0 and derived allele as 1. At each SNP site, all reference haplotypes and the target form a coalescent tree with *n* + 1 leaves. This tree is unknown but represents the complete information that imputation can possibly use (Figure 1). If the target’s most recent common ancestor (MRCA) with the reference sample occurs on a branch that is ancestral to *u* ≥ 1 reference haplotypes, all *u* present day descendants of that branch are the most closely related reference haplotypes. We assume that the imputation algorithm identifies these most closely related reference haplotypes, then assigns the mean genotype of them to the target haplotype as the imputation dosage.

Under this assumption, we consider three scenarios: (1) The mutation generating the target variant is ancestral to the time to the most recent common ancestor (TMRCA) of the target haplotype and all its most closely related reference haplotypes. In this case, it will be imputed correctly with dosage 1 (Figure 2 a). (2) The mutation occurred more recently than the TMRCA and the mutation occurred on the branch to the target haplotype. Then the target variant is not polymorphic in the reference sample and will always be falsely imputed to be the ancestral allele (Figure 2 b). (3) The mutation occurred more recently than the TMRCA and the mutation occurred on the branches to the reference haplotype. Then some or all of the most closely related reference haplotypes carry the derived allele while the template haplotype carries the ancestral allele and the reference sample will be imputed to carry the derived allele with some dosage> 0 (Figure 2 c and d).

**Figure 2:**
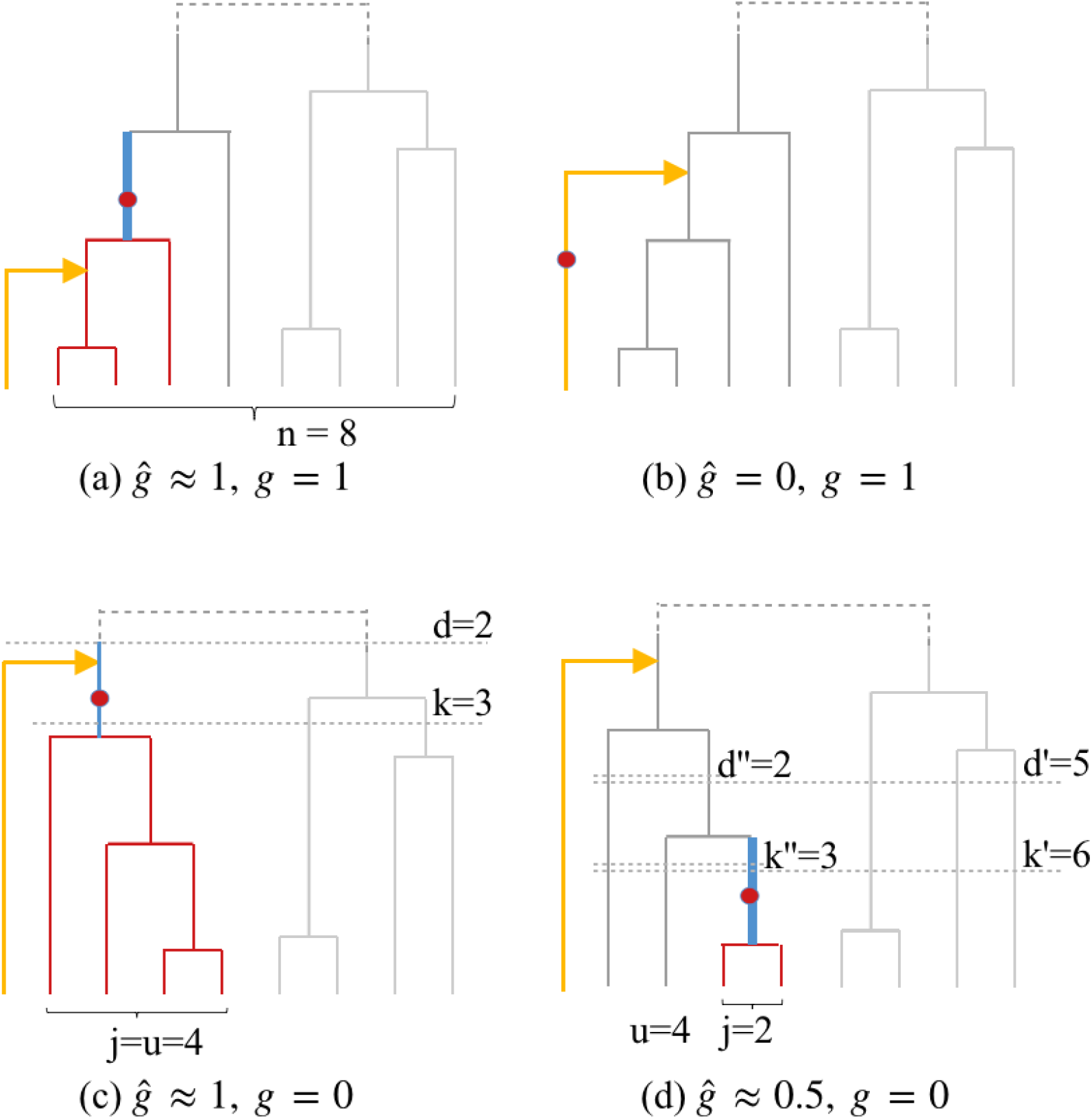
Possible scenarios when imputing one target haplotype (yellow lineage) using a reference panel with *n* haplotypes. Red lines are haplotypes carrying a mutation that occurs on the blue branch. Gray horizontal dashed line indicate the time intervals where an event of interest occurs. a) The derived allele arises ancestral to the MRCA of the target with the reference haplotypes and is shared between the target (*g* = 1) and all most closely related haplotypes. Thus it will be correctly imputed (*ĝ* ≈ 1). b) The derived allele arises on the branch of the target (*g* = 1) and is thus absent from the reference. It will be falsely imputed to be ancestral (*ĝ* = 0). c) The derived allele arises after the MRCA of the closest reference haplotypes before coalescing with the target haplotype. The target haplotype does not carry the derived allele (*g* = 0), but all closest reference haplotypes do. The target will be imputed to carry the derived allele (*ĝ* ≈ 1). d) The derived allele arises on a branch ancestral to *j* = 2 of the closest reference haplotypes before the MRCA of all *u* = 4 closest haplotypes. The target haplotype does not carry the derived allele (*g* = 0). The target’s derived allele dosage will be imputed to reflect that *j* of *u* closest haplotypes carry the derived allele (*ĝ* ≈ *j/u* = 0.5).

We focus on the third scenario where *j* of the *u* most closely related templates carry the derived allele while the target does not. The Li and Stephens model assumes a single closest haplotype, and in the HMM implementation the imputed fractional genotype (dosage) is a weighted average of multiple reference haplotypes based on their posterior probabilities of being that right template. When the *u* are equivalently close to the target, they are expected to have equal posterior probabilities. Thus the imputed dosage *ĝ* is *ĝ* = *j*/*u* ∈ (0, 1] while the true genotype *g* is *g* = 0 (“false positive”), resulting in loss of information in downstream analysis.

In the following sections, we derive the probability for all possible (*j, u*) configuration, conditional on observing the derived allele count *j* in the whole reference. For simplicity, we distinguish the one target haplotype from the rest *n* reference ones, making the whole tree size *n* + 1, although they are exchangeable under the assumption of homogeneity and random mating. We will always consider time backward with *t* = 0 being the current generation.

### 2.1 Number of most closely related templates

We first give the probability of having *u* equally good templates at any random position for a target haplotype: *P*(*u*; *n*). This probability depends only the topology of the genealogy, independent from mutation events. We leverage that the probability that a set of *k* lines coalesce before they coalesce with any line among the rest *n* − *k* [11] is

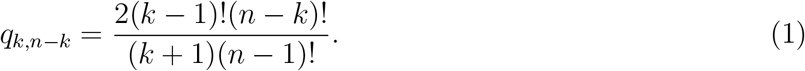

The generative process can then be imagined as three steps: *u* templates coalesce first before their MRCA meets the target; then the resulting branch of size *u* + 1 meets the rest of the tree. Finally we sum over all possible sets of *u* templates.

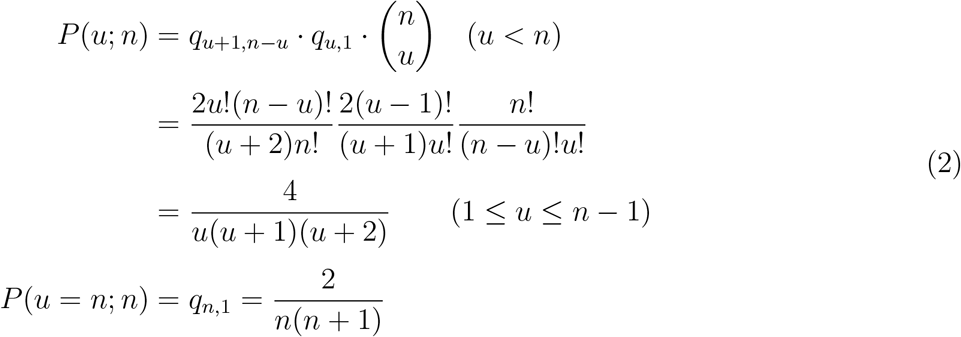

The expected number of best templates is close to 2 when the reference is large:

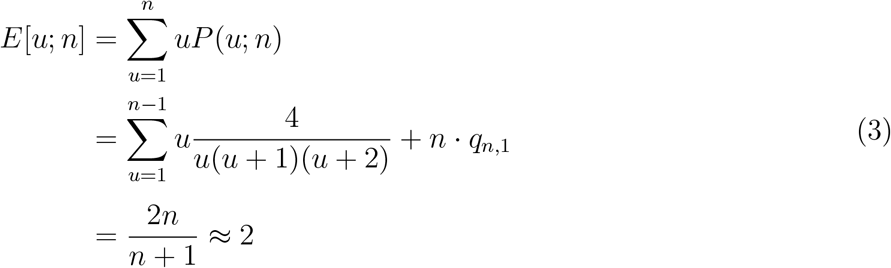

### 2.2 High certainty error

We now derive the probability of imputing the target to carry the mutation with dosage *ĝ* = 1 while the truth is *g* = 0, conditional on the observed derived allele count in the reference panel: *P*(*ĝ* = 1, *g* = 0|*j*; *n*). This happens when the target haplotype first coalesces with a branch (*l*) of size *j* carrying a mutation (Figure 2 c).

Throughout the following derivation, we use the number of ancestral lines *k* for the current day sample to keep track of coalescent time and to connect topology and branch lengths. We first introduce some quantities useful for our derivation:

Let *P*(*j, k*; *n*) be the probability for *j* lines to reach their MRCA at the coalescent event that reduces the overall number of ancestral lines from *k* + 1 to *k*, without any of the *j* lines coalescing with any of the *n* − *j* other lines first. Let *P*_0_(*n, d*) be the probability for one line to encounter no coalescent event for (at least) the first *n* − *d* events. Let *P*(*m* ≥ 1|*l*) be the probability of having at least one mutation event on a branch with length *l*, and let *P*(*L* = *l*|*k, d*; *n*) be the probability density function for the length of one internal branch starting when there are *k* lines left and ending when there are *d* lines left, in a tree of size *n*. Using these terms, we can now calculate

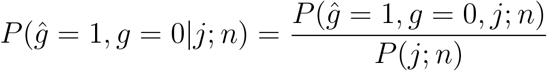

We calculate the joint probability *P*(*ĝ* = 1, *g* = 0, *j*; *n*) in the following three steps by conditioning on *d* and *k* (Eq. 4, Figure 2 c). We calculate *P*(*j*; *n*) by adapting the second step (Eq. 5).

1. To coalesce in the (*n* − *d* + 1)-th coalescent event, when *d* lines remain on the tree, the target haplotype cannot coalesce in the first *n* − *d* events. By definition of *P*_0_, this is *P*_0_(*n* + 1, *d* + 1).
2. A branch of size *j* ancestral to all most closely related reference haplotypes arises in the reference at the (*n* − *k*)-th event (*P*(*j, k*; *n*)) and encounters a mutation before it coalesces with the target branch in event *n* − *d* + 1 (*g*(*k, d*; *n*)). Here the probability for a mutation to occur on that particular branch is computed by integrating over all possible branch length *l* given *k* and *d*: 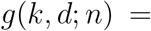 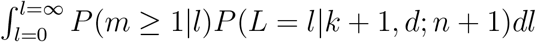.
3. The size-j branch does not coalesce till the (*n* − *d*)-th event and then coalesces with the target: 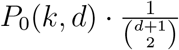.

Multiplying the probabilities of these sequential events and summing over all possible values of *d*, *k* give the joint probability:

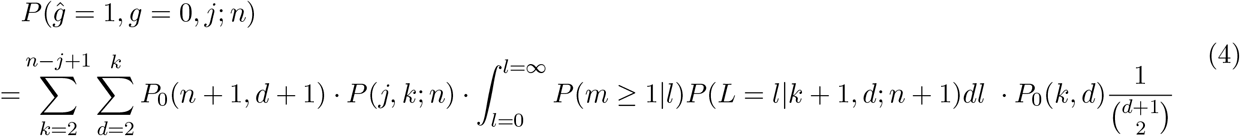

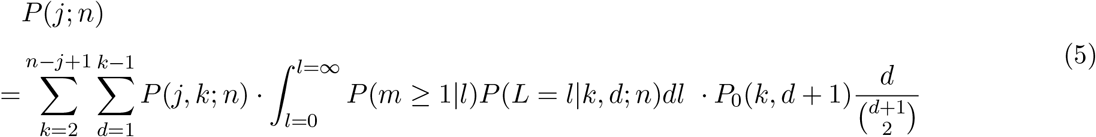

To compute *P*(*j, k*; *n*), define 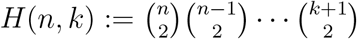, the number of possible configurations of the topological history for *n* lines to coalesce till there are *k* ancestral lines left. Then

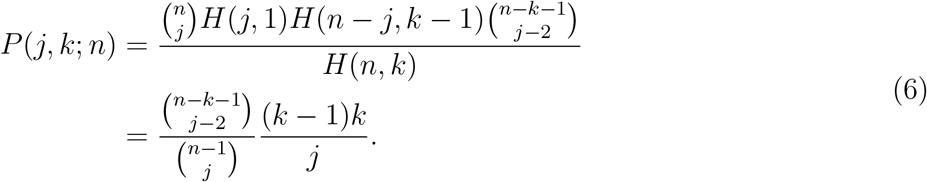

Next we consider the integral corresponding to the mutation event *g*(*k, d*; *n*) (Eq. 7). Let *E*[*T_i_*; *n*] denote the expected time for the number of ancestral lines to go from *i* to *i* − 1 in the coalescence process for a sample of *n* current haplotypes, which depends on the population history model. Thus, *g*(*k, d*; *n*) depends on mutation rate *μ* and expected coalescent time intervals *E*[*T_k_*; *n*], which can be numerically computed [12] or approximated by Monte Carlo.

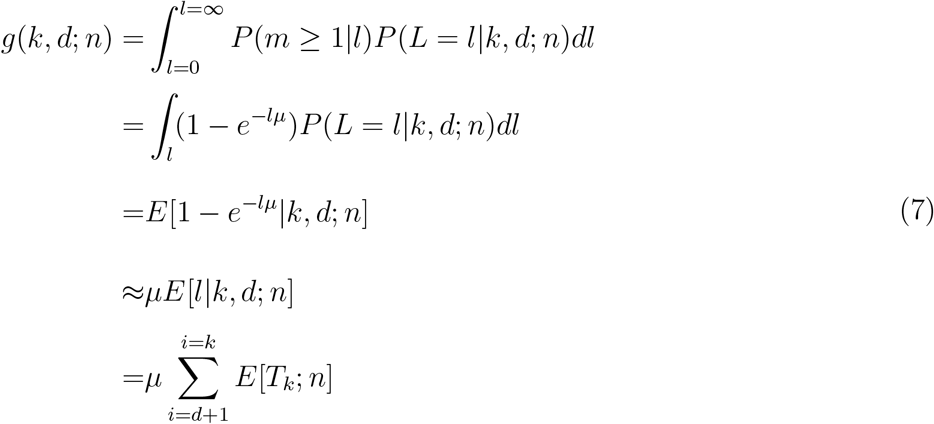

Substituting Eq. 6,7 and 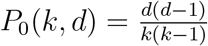 into Eq. 4,7, we now have the conditional probability fully defined:

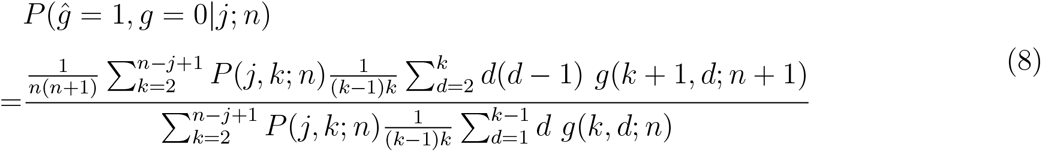

Following similar logic as above, we then consider the conditional probability distribution for fractional dosages. We derive *P*(*ĝ* = *j/u, g* = 0|*j*; *n*), the probability of having *u* equally closely related best templates, *j* of which carry the mutation, for a target actually carrying the ancestral allele, conditional on observing the derived allele count as *j* in a reference of size *n* (figure 2 d). Details of derivation are in the Appendix A.

### 2.3 Modelling r^2^ as a function of misidentification proportion

The above derivation fully characterizes the distribution of the imputed dosages assuming we perfectly identify the optimal templates. While it is difficult to model all the possible sources of error in the real imputation process, we can use the total weight (*q*) attributed to haplotypes outside the set of most closely related templates to measure the identification error. Here we derive its influence on the square of correlation coefficient (*r*^2^) between the true genotype and imputed dosage.

Let *g* be the true genotype, 1 for the derived allele and 0 for the ancestral allele; let *f* be the derived allele frequency, so *E*[*g*] = *f*. Let 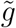 and *ĝ* be the imputed dosage with and without misidentification respectively.

Let *z* be the random variable representing the proportion of carriers of the minor allele among the sub-optimal templates contributing to the imputed dosage, so that 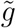 can be modeled as a mixture 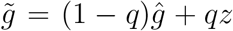. If we assume that the sub-optimal templates are sampled at random from the population and each of the suboptimal templates is given equal weight, *z* is the average genotype of *m* random haplotypes, so *E*(*z*) = *f* and *V*(*z*) = *V*(*g*)/*m* = *f*(1 − *f*)/*m*.

When the optimal templates for a target sample provide the perfect information (*ĝ* = *g*), misidentified templates may be the alternate allele and therefore increase imputation error. However, where the target sample carries the ancestral allele but some of its optimal templates carry the derived allele, sub-optimal templates attenuate the resulting error. Nevertheless, here we show that in expectation imputation error will always reduce the correlation between the true and the imputed genotype. Let’s express 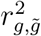 in terms of quantities regarding the theoretical imputed dosage *ĝ*.

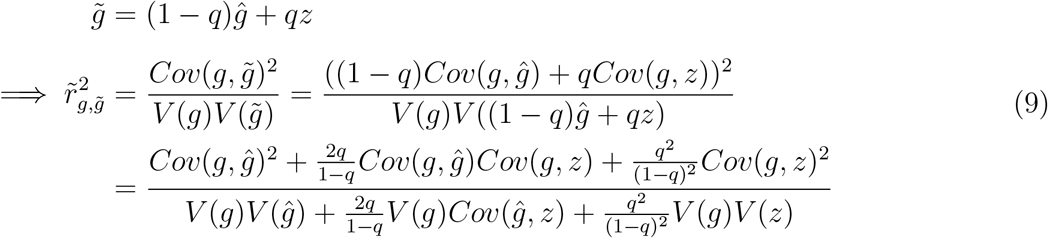

Comparing to the theoretical 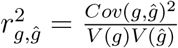:

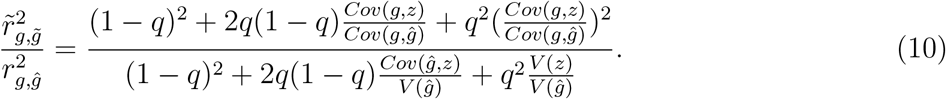

Since *ĝ* is based on a few reference haplotypes and equal to *g* with large probability, we can reasonably assume *Cov*(*g, z*) ≈ *Cov*(*ĝ*, *z*), *Cov*(*g, ĝ*) ≈ *V*(*g*) and *Cov*(*g, z*) < *Cov*(*g, ĝ*) (misidentified haplotypes are more distant). Therefore the middle terms in the numerator and denominator of Eq. 10 are approximately equal; the difference between the *r*^2^’s is governed by the last terms, which have a ratio strictly smaller than one: 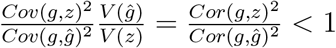.

If misidentified haplotypes are close to random draws from all reference haplotypes (*Cov*(*g, z*) ≈ 0), the ratio simplifies to

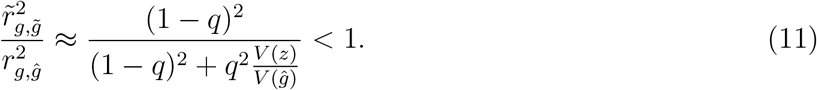

In practice, the sub-optimal templates contributing to the imputed dosage are more likely to be from lineages closer to the target, so their effect on imputation accuracy would be smaller than that from the above assumptions.

### 2.4 Length of haplotype before the next recombination breakpoint

Next we aim to derive the distance between consecutive recombination events on an external branch. For the purpose of modelling imputation, these distances represent the lengths of segments that are copied from the same haplotype. Let *X* be the genetic distance to the next observed recombination event on the target, and *T* be the length of its corresponding external branch. We calculate *f_X_*(*x*) = ∫ *f*_*X*|*T*_ (*x*)*f_T_*(*t*)*dt* (we will use *f*(·) for continuous and *P*(·) for discrete distributions). Similar to above, we compute the probability density function of *T* by considering it first coalesces at the (*n* + 1 − *k*)th event. *T* |*k* is the sum of *n* + 1 − *k* coalescent time intervals, thus following a convolution distribution as the sum of *n* + 1 − *k* exponential with different rates. Conditional on *T*, *X* follows an exponential distribution with rate 2*T* since a recombination event on either the current template or the target haplotype results in a switch of template.

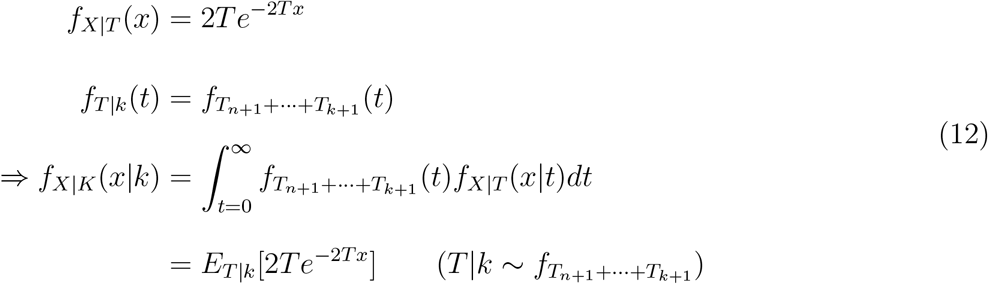

When population size is constant, 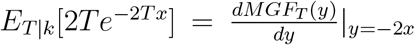, where *MGF_T_* s the moment generating function of the hypoexponential distribution *T*|*k*. When population size changes through time so time intervals are correlated, we can approximate the expected value with 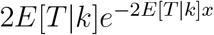 and compute *E*[*T* |*k*] with Monte Carlo.

The probability for a haplotype to first coalesce at the (*n* + 1 − *k*)th event is 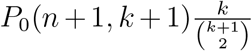, with *P*_0_(*n* + 1, *k* + 1) being the probability of not coalescing with any lineage in the first (*n* − *k*) events (starting from total sample size *n* + 1). Therefore we have the distribution of *X* is

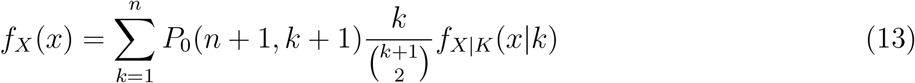

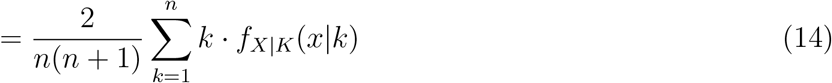

### 2.5 Simulation

We performed a standard Kingman coalescent simulation under population history models suggested by recent studies (detailed in the Results). We simulate size-(*n* + 1) trees with *n* varying from 500 to 50*k*. For each generated tree, we keep track for all subtree configurations (1 + *u*(*j*)) (one external branch first coalesces with a branch of size *u*, which contains a branch of size *j*) of interests, and record the length of this size *j* branch. The sum of all such relevant branch lengths in the tree divided by the number of present haplotypes *n* + 1 is one realization of the joint probability *P*(*ĝ* = *j/u, g* = 0, *j*; *n*), which will then be approximated by averaging such realizations over all simulated trees. To approximate the probability of observing *j* derived alleles in the reference panel, we simulated another independent set of trees of size *n* and summed over the lengths of all size *j* branches to get sample frequency spectrum. We present results from 50*k* independent simulations under each parameter setting throughout this article.

### 2.6 Empirical Evaluation

To evaluate the predictions of our model, we imputed genotypes from the Michigan Genomics Initiative (MGI) [13] using different imputation panels. To assess imputation quality, we masked some genotypes and compared the imputed genotypes to the genotypes generated by the array. MGI is collected from the patient population of the University of Michigan Hospitals and thus mostly (> 90%) consists of individuals of European decent. The MGI individuals analyzed here are genotyped on a Illumina Infinium CoreExome chip with ~ 60, 000 custom markers, providing sufficient low frequency variants to assess imputation performance for rare variants.

We considered five reference panels: (1) The Haplotype Reference Consortium (HRC) r1.1 reference panel (*n* = 32, 470 individuals of mostly European descent, ≈ 39*M* variants) [14], (2) the 1000 Genomes Project (1KGP) variant calls from low-coverage sequencing (*n* = 2, 548, ≈ 78*M* variants) [15], (3) the 1KGP variant calls from high-coverage sequencing (*n* = 2, 504, ≈ 100*M* variants) (Michael Zody, personal communication), (4, 5) subsets of the high- and low-coverage 1KGP variant calls containing only samples from the European super population (*n* = 503). Note that the HRC reference panel only provides varaints with minor allele count ≥ 5 [14]. We subsetted all 1KGP variant reference panels to the set of overlapping individuals and markers and filtered out monomorphic and singleton sites, resulting in 2, 503 samples and 41*M* varaints for panels 2 & 3 and 503 samples and ≈ 13*M* variants for panels 4 & 5. We also re-phased all 1KGP panels using Eagle (v2.4.1) [16] without a reference panel to avoid confounding due to differing phasing quality.

Using each reference panel, we imputed all autosomal variants by running Minimac4 (v1.0.0) [17] on a computing cluster maintained by the Center for Statistical Genetics at the University of Michigan in Ann Arbor. We set Minimac4 parameters to use 16 cpus and output data in genotype, estimated alternate allele dosage, and estimated haploid alternate allele dosage formats. For all rare variants genotyped in MGI, we estimated imputation quality using the squared Pearson correlation coefficient between known and imputed genotypes.

## Data Availability

1000 Genomes low coverage sequence data were downloaded from ftp://ftp.1000genomes.ebi.ac.uk/vol1/ftp/data_collections/1000_genomes_project/release/20190312_biallelic_SNV_and_INDEL/. 1000 Genomes were downloaded from ftp://ftp.1000genomes.ebi.ac.uk/vol1/ftp/data_collections/1000G_2504_high_coverage/.

## Software Availability

Codes used to perform analytical calculation and coalescent simulation are available from https://github.com/Yichen-Si/ImputationBound.

## 3 Results

We derived an analytical approach to calculate the impact of model misspecification of the current imputation framework with our coalescent model on rare variants (see the Method). Assuming a Li and Stephens model[6] based imputation algorithm correctly identifies all and only those haplotypes in the reference that are most closely related to the target sample (”closest templates”), we calculated the theoretical error rate as a function of reference size *n* and the derived allele count (DAC). Using coalescent simulations, we confirm the analytical results, and incorporate general population models. We compared our theoretical predictions with empirical imputation accuracy observed in the Michigan Genomics Initiative (MGI), with the 1000 Genomes[18] or the Haplotye Reference Consortium[19] as reference.

### 3.1 The number of closest template haplotypes

In the coalescence context, identifying a single reference haplotype that is most similar to the target haplotype is equivalent to selecting the reference haplotype that has the MRCA with the target haplotype or, equivalently, whose lineage is the first to coalesce with the target. However, by the time this lineage coalesces with the target, it may be ancestral to multiple reference haplotypes (Figure 1). In this case, those reference haplotypes are equally closely related with the target, each of them in expectation providing the same amount of information for imputing the target sample. Thus, the model assuming one single best template is misspecified, and imputation is more likely to be ambiguous when the number of closest templates is larger.

The probability for the target haplotype to first coalesce with a lineage having *u* descendants in the reference is

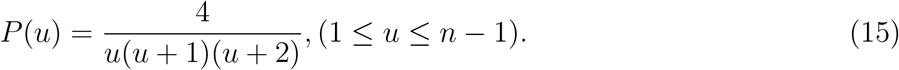

Note that the derivations of this probability do not condition on the presence of a variable site. Thus,this probability depends only on the topology of the coalescent tree, independent of population history (see the Method). Interestingly, the probability distribution of the number of best matches does not depend on the size of the reference panel (*n*), except for the extreme case *u* = *n*, where the entire reference has equal genetic distance to the target. In that case, 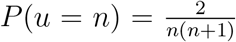. The expected number of closest templates 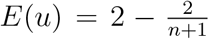, approaching 2 as the size of the reference panel increases.

From equation (15), we see that with probability 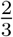, the target haplotype first coalesces with an external branch (*u* = 1) and the reference contains exactly one best match. Thus, the Li and Stephens model of exactly one best template is misspecified with probability 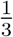. The probability of *u* equally good templates drops rapidly with increasing *u* (Table 2). With probability 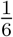, the target haplotype first coalesces with an internal branch with two descendants in the reference (*u* = 2), while the probability is 0.0108 that the target haplotype first coalesces with an internal branch with more than 10 descendants in the reference (*u* > 10).

**Table 1:**
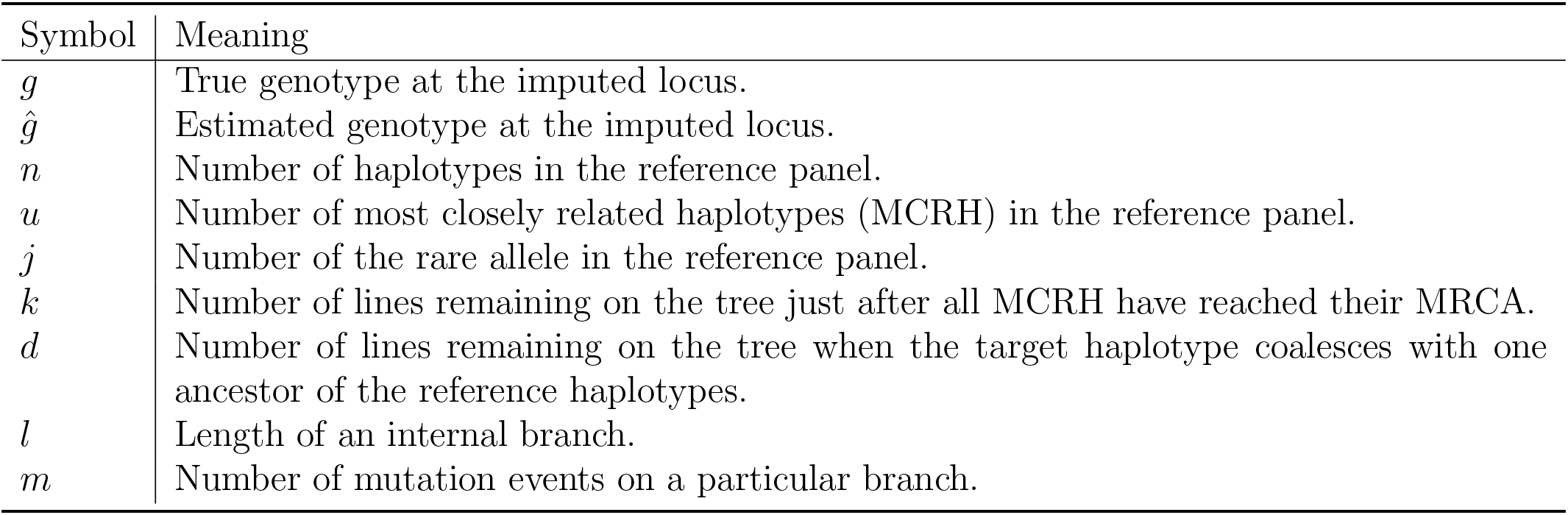
Notation for key quantities

| Symbol | Meaning |
| --- | --- |
| $g$ | True genotype at the imputed locus. |
| $\hat{g}$ | Estimated genotype at the imputed locus. |
| $n$ | Number of haplotypes in the reference panel. |
| $u$ | Number of most closely related haplotypes (MCRH) in the reference panel. |
| $j$ | Number of the rare allele in the reference panel. |
| $k$ | Number of lines remaining on the tree just after all MCRH have reached their MRCA. |
| $d$ | Number of lines remaining on the tree when the target haplotype coalesces with one ancestor of the reference haplotypes. |
| $l$ | Length of an internal branch. |
| $m$ | Number of mutation events on a particular branch. |

**Table 2:** Probability of multiple reference haplotypes (“No. Templates”) being most closely related to the target haplotype.

| No. Templates | 1 | 2 | 3 | 4 | 5 | 6 | 7 | 8 | 9 | 10 |
| --- | --- | --- | --- | --- | --- | --- | --- | --- | --- | --- |
| Prob. | 0.6667 | 0.1667 | 0.0667 | 0.0333 | 0.0190 | 0.0119 | 0.0079 | 0.0056 | 0.0040 | 0.0030 |

### 3.2 Impact on imputation accuracy

As we have just demonstrated, the model assumption that each target haplotype has a single most closely related template haplotype is a model misspecification for 1/3 of the genome where there are multiple closest templates. This model misspecification contributes to imputation error beyond the typical source of error created by failing to identify the most closely related haplotype. To isolate this additional error, we now evaluate the imputation of a missing genotype assuming that all closest templates are correctly identified but analyzed under a model that assumes a single best haplotype. In this scenario, imputation algorithms will correctly impute all non-singleton variants if the target haplotype has a single most closely related template haplotype. Imputation errors can only occur if the target haplotype has more than one closest template, i.e. a model misspecification. In this case, imputation algorithms consider all these template haplotypes to be equally likely to be the “best” template. Accordingly, it will interpret identifying multiple equally close templates as uncertainty in identifying the best template. For variants that differ between these template haplotypes, the imputed genotype is then usually the average of the DAC of those templates, a fractional genotype (dosage). Thus, for one given target haplotype carrying the ancestral allele to be imputed with a non-zero DAC dosage (“false positive”), some of its templates have to carry the derived allele.

While the probability of this misspecification is independent of population history, the probability of a mutation event causing an imputation error depends on the history of the population template and target haplotypes are sampled from. In our primary analysis we consider a population history model approximating European population history [20, 21]. Starting from an ancestral population with effective population size 10^4^, it undergoes a bottleneck with *N_e_* = 2 · 10^3^ for 100k years (approximately 3450 generations), then grows with an accelerated rate (faster than exponential growth [22]) to *N_e_* = 10^7^ during the most recent 10k years. The effect of these model parameters on the following results is marginal within a model space reasonable for human population (Appendix B).

We first consider the special case where all the closest templates for a target carry the derived allele, while the target haplotype carries the ancestral allele. In this case, the imputed dosage of the derived allele is 1 when the truth is 0 (Figure 2 a). By integrating over all possible tree shapes and branching times (see the Method), we calculated the probability for this configuration for reference sample sizes of 500, 5000, 20,000 and 50,000 individuals. We verified all results using computer simulations. Across all considered reference sizes, the probability of this completely wrong imputation for one imputed haplotype is small ranging from 10^−3^ to < 10^−6^ (Figure 3 a, Table 3). Holding reference size constant, this probability decreases with rising DAC. Similarly, holding DAC constant, this probability of imputation error decreases with increasing number of templates.

**Figure 3:**
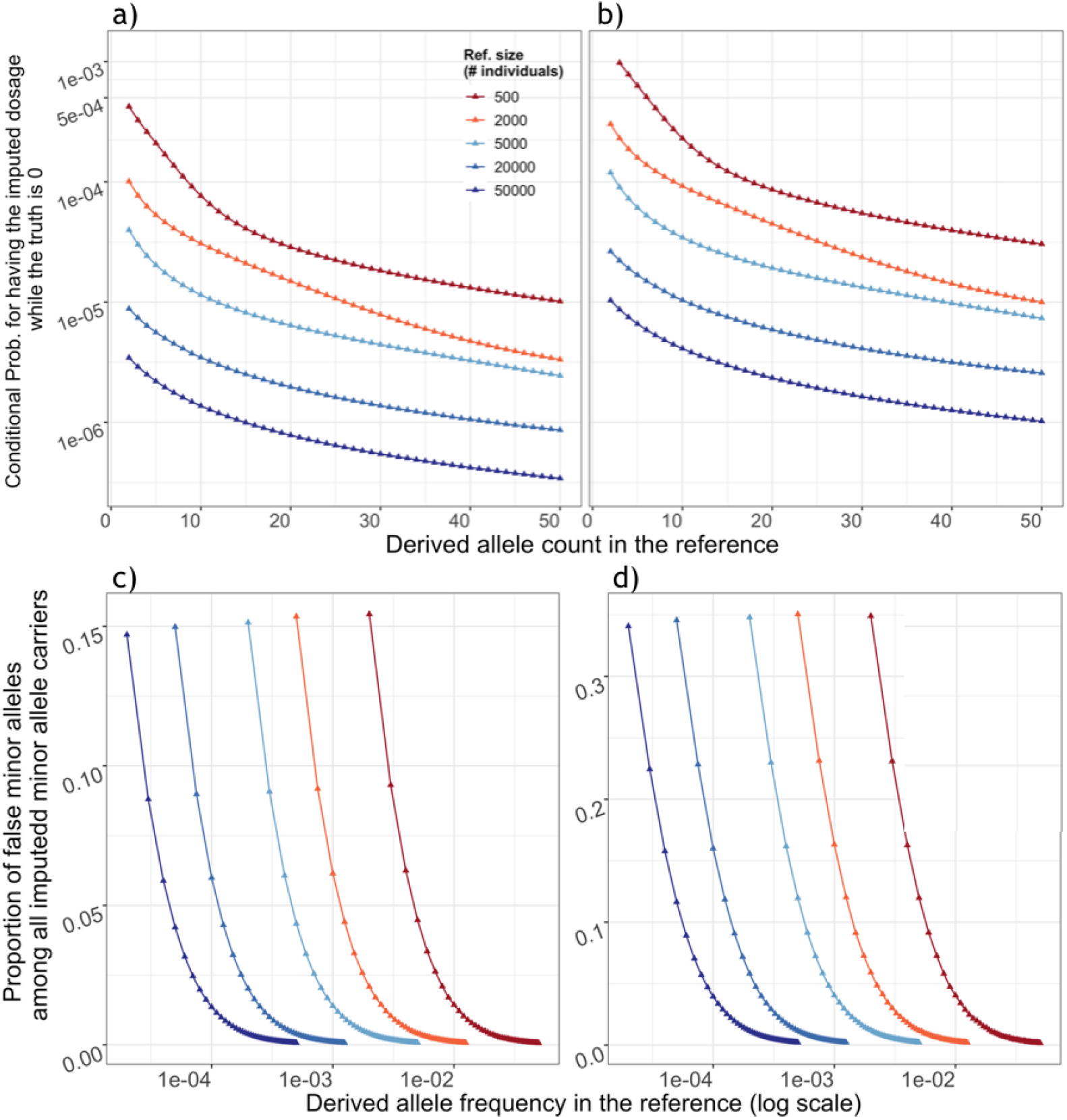
Upper panel: False discovery rate among non-carriers. Given a target haplotype carrying the ancestral allele (true dosage 0), the probability (y-axis, in log scale) of having a) all the closest templates in the reference panel carrying the derived allele thus an estimated dosage 1; or b) having non-zero estimated dosage for the derived allele. Lower panel: proportion of false positives among all imputed carriers. For a target sample, the proportion (y-axis) of haplotypes with the ancestral allele among c) individuals with estimated dosage 1 or d) haplotypes with non-zero dosage. Both results are conditional on the DAF in the reference (x-axis); color indicates the size of reference panel, in the number of individuals. Results for reference size below 20,000 are from analytical calculation while those for 20,000 and 50,000 are from coalescence simulations.

**Table 3:**
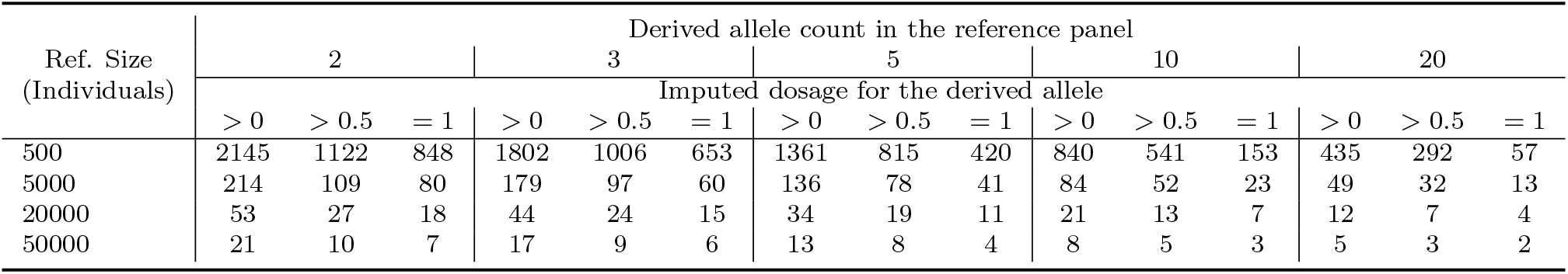
Expected number of imputation errors. Consider imputing a sample dataset with one million individuals using a reference panel containing 500 to 50000 individuals from the same population as the sample, the expected number of individuals who are homozygous for the ancestral allele (*g* = 0) but have an imputed genotype dosage *ĝ* as *ĝ* > 0, *ĝ* > 0.5 or *ĝ* = 1. Each combined column represents one DAC in the reference. Results are from 50*k* independent coalescence simulations, each data point is an average of over 10^7^ loci.

To put these error rates in relationship with the number of true carriers of the derived allele, we assume that the derived allele frequency (DAF) in the target population is the same as that in the reference population. For instance, consider a site with two copies of the derived allele in a reference of 500 individuals (MAF = 0.002). For this site, we expect to impute one individual as false carrier for every 1210 individuals (Figure 3). Among these 1210 individuals, we also expect 4.84 true carriers of this variant assuming the same MAF in the target population. Thus, for variants that are doubletons in a reference sample of 500 individuals, about one in six imputed rare alleles will be a false positive (Table 3). More generally, for a fixed DAC in the reference, the probability of falsely imputing the derived allele decreases with increasing reference size while the number of true carriers of the derived allele also decreases. As a result, the proportion of falsely imputed derived alleles decreases only moderately with increasing reference size. For example, for variants that are doubletons in the reference, the proportion of false positive derived alleles decreases only from 17.5% for a reference size of 500 to 14.7% for a reference size of 50,000.

If we now consider the more general case where at least one of the closest templates carry the derived allele while the target carries the ancestral allele (dosage 0). In this case the imputed dosage *ĝ* > 0 (Figure 3). Across reference sizes of 500, 5000, 20,000 and 50,000 individuals, we calculated the probability of *ĝ* > 0 for each imputed haplotype. This probability is notably larger than the probability of falsely imputing dosage 1, ranging from ~ 10^−3^ to ~ 10^−5^ for doubletons (with DAC 2, Figure 3 b). This probability decreases with rising DAC and increasing reference size.

From these probabilities, we calculate the expected number of haplotypes carrying the ancestral allele (*g* = 0) that are falsely imputed to have either a non-zero derived allele dosage (*ĝ* > 0), a higher dosage *ĝ* > 0.5 or a dosage of 1 for every million target individuals (Table 3). Haplotypes with *ĝ* > 0.5 represent cases where a “best-guess” imputation algorithm would infer the alternate allele. Beyond the previously described impact of DAC and reference size, these results show that for DAC < 5, about half of all falsely imputed derived alleles have a dosage > 0.5. This observation can be explained by the fact that for a given DAC, observing a dosage < 0.5 requires a larger number of equally good templates than observing a larger dosage. As large numbers of equally good templates are rare (Table 2), the proportion of higher dosage among all falsely imputed non-zero dosages increases with the DAC.

To summarize the impact of this error on association tests, we calculated the squared correlation coefficient (*r*^2^) between the imputed dosages and the true genotypes using simulations, both under the assumption that model misspecification is the only source of error and under the assumption that some reference haplotypes are falsely identified as being most closely related. *r*^2^ is a commonly used measure for imputation quality, as it is directly related to statistical power in downstream association tests [23, 24, 25].

Still assuming that closest templates are identified perfectly, we generated the distribution of expected *r*^2^ for a range of minor allele counts (MAC) and reference sizes using simulations (see the Method). The expected *r*^2^ increases monotonically with DAF in the reference and reference size (Figure 4). If we consider the increase with DAF across reference sizes, we observe that the curves of expected *r*^2^ values look very similar for each reference panel size, only shifted by a factor of 1/panel size. In other words, *r*^2^ of variants with the same MAC remains almost the same across all considered reference sizes. For example, the average *r*^2^ among doubletons is 0.822 in a reference size of 500 and 0.831 in a reference sample size of 50,000, where its frequency is only 1/100 of the former. For variants observed ten or more times in the reference, the expected *r*^2^ is > 0.97 regardless of the size of the reference, as model misspecification do not play a mayor role for the imputation quality of these variants, but for variants observed less often, *r*^2^ decreases rapidly with decreasing allele count. To include other sources of error in the imputation process, we model the false identification of reference haplotypes that are more distantly related to the target as templates. Those distantly related haplotypes are less likely to carry the same allele as the target haplotype and thus introduce an additional source of error. We parameterize this identification error as the sum of all probabilities assigned to falsely identified templates, denoted as *q* (see the Method). In practice, this imputation error depends on the choice of imputation algorithm, marker density, quality of genotyping and statistical phasing; detailed modeling of these factors is beyond the scope of this paper.

**Figure 4:**
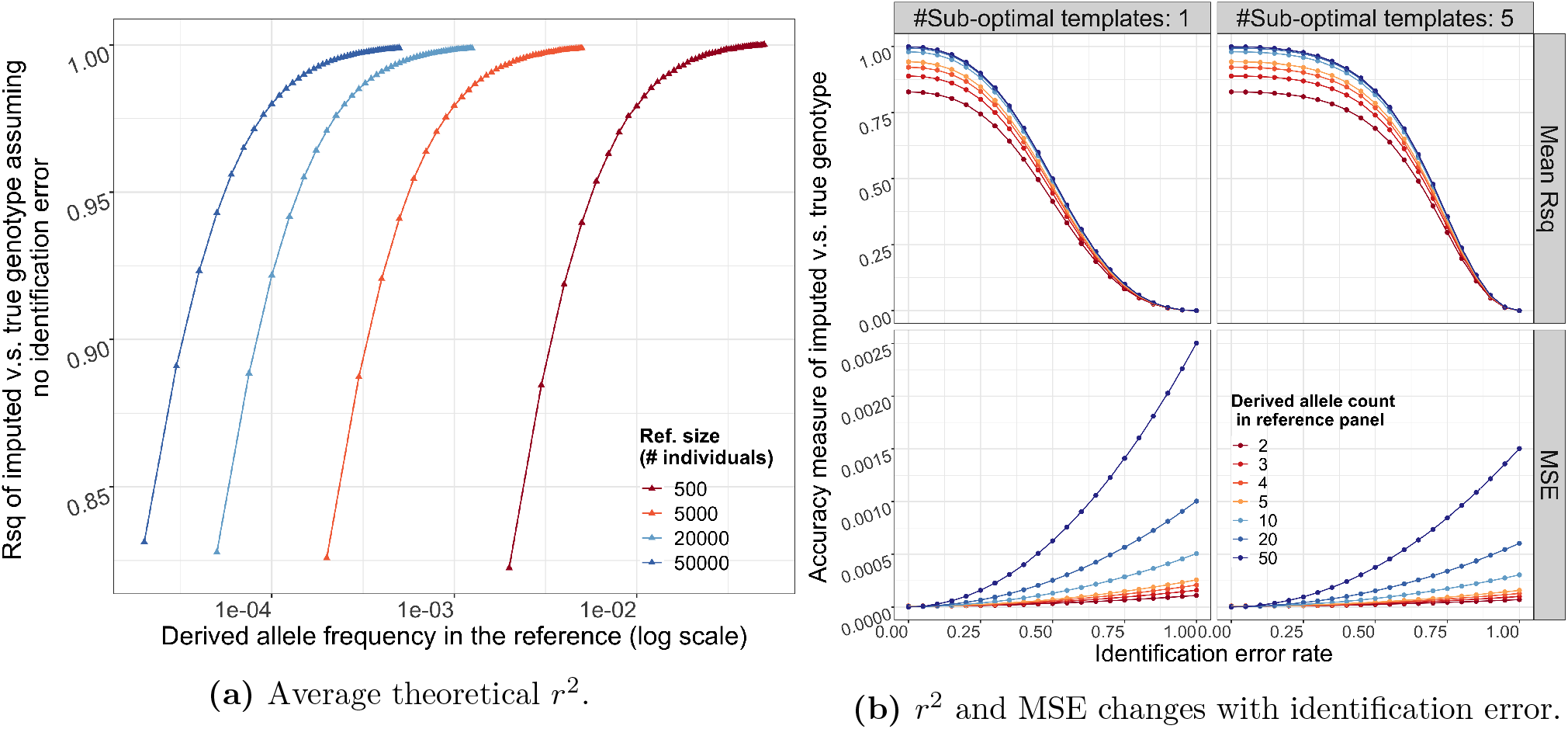
a) The upper bound of squared correlation (*r*^2^) (y-axis) between imputed and true genotype dosages, conditional on the DAF in the reference (x-axis). Color indicates the size of reference panel. b) The *r*^2^ (top) and MSE (bottom) when a certain proportion of weight (x-axis) is attributed to sub-optimal templates. The impact depends on the absolute number of sub-optimal templates, we show examples of 1, 5 in the left and right columns. Here we fix the reference size to 20*k*. Results are from 50*k* independent coalescence simulations, each data point is an average of over 10^7^ loci.

As expected, imputation error of the variant increases with increasing *q* shown as the mean squared error (MSE) of the imputed genotype (Figure 4b). As the imputed genotype for all individuals converges to the allele frequency *f* as *q* goes to 1, the MSE increases faster with higher *f*. Similarly, the *r*^2^ between the true genotype and the imputed genotype decreases with increasing *q*. However, including some suboptimal haplotypes only has a small effect for rare variants. If we assume a single sub-optimal template, *q* < 10% only has marginal effect on *r*^2^, when *q* increases beyond this threshold the decrease of *r*^2^ becomes almost linear. This threshold depends on the number of sub-optimal templates: with a larger number of templates, the variance of the imputed genotype decreases and the effect of including suboptimal templates is more deterministic (see the Method). Accordingly, if the number of suboptimal templates is larger, the threshold where *r*^2^ starts to decrease with *q* is larger while the MSE is lower (Figure 4b). For example, consider imputing a variant that is a doubleton in the reference with 20k haplotypes. Assuming no identification error, 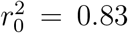. If we identify one sub-optimal templates with posterior probability 0.3 as well as all the closest templates, the expected *r*^2^ is 0.70 and the MSE is 1.80 × 10^−5^. If we instead identify 5 sub-optimal templates with total posterior probability 0.3 as well as all the closest templates, the expected *r*^2^ increases to 0.80 while the MSE decreases to 1.44 × 10^−5^.

For variants with DAC < 5 in the reference, the error caused by model misspecification dominates for a wide range of *q*; while for variants of DAC over 20 the error caused by model misspecification is negligible and the 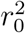 with perfectly identified optimal templates is above 0.99 regardless of the reference size (Table 4), thus the identification error dominates the empirical imputation error.

**Table 4:** The maximal squared correlation can be achieved. Numbers in the parentheses are standard deviation summarized over 10^7^ loci, representing the variation among variants in a population sample. Results are from 50*k* independent coalescence simulations.

| Ref. Size<br>(Individuals) | Derived allele count in the reference |  |  |  |  |  |
| --- | --- | --- | --- | --- | --- | --- |
|  | 2 | 3 | 4 | 5 | 10 | 20 |
| 500 | 0.822(0.089) | 0.885(0.067) | 0.919(0.051) | 0.940(0.041) | 0.979(0.017) | 0.995(0.0055) |
| 2000 | 0.823(0.089) | 0.886(0.066) | 0.920(0.051) | 0.940(0.040) | 0.979(0.016) | 0.994(0.0054) |
| 5000 | 0.826(0.088) | 0.887(0.065) | 0.921(0.050) | 0.941(0.040) | 0.979(0.016) | 0.994(0.0055) |
| 20000 | 0.828(0.087) | 0.889(0.065) | 0.922(0.050) | 0.942(0.040) | 0.980(0.016) | 0.994(0.0053) |
| 50000 | 0.831(0.087) | 0.891(0.065) | 0.923(0.050) | 0.943(0.039) | 0.980(0.016) | 0.994(0.0053) |

### 3.3 Comparison with empirical imputation accuracy

We imputed genotypes of 56, 984 participants in Michigan Genomics Initiative (MGI) using two standard reference panels, the 1000 Genomes (1KG, 2504 individuals) and the HRC (32, 470 individuals). For variants present in the MGI dataset, we calculated *r*^2^ between observed genotypes and imputed genotypes and stratified these by the MAC of the variant in the reference panel (figure 5).

**Figure 5:**
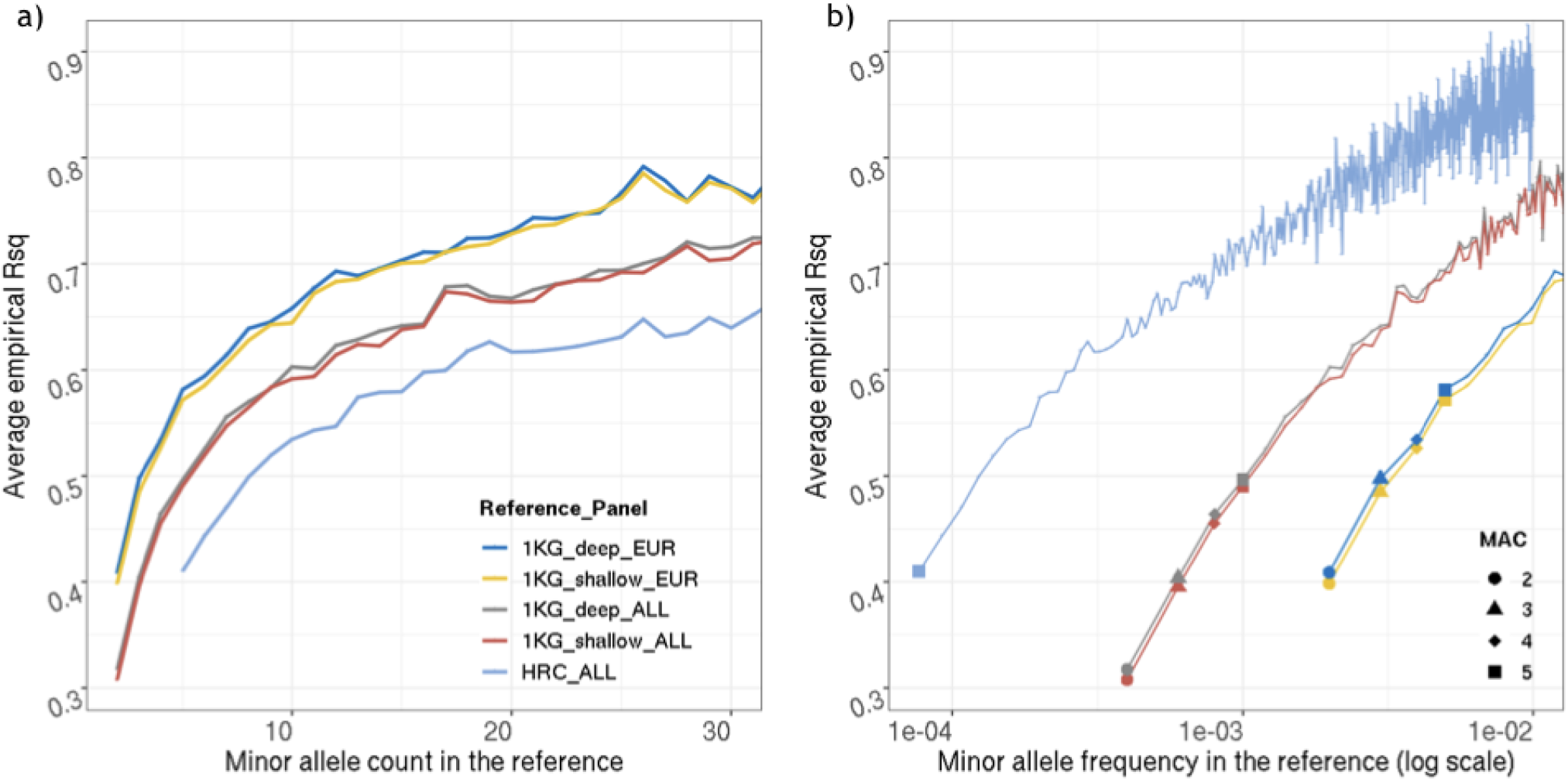
Imputation accuracy empirically evaluated in MGI using different reference panels. Y-axis is average *r*^2^ evaluated at genotyped sites, x-axis is a) MAC or b) MAF in the reference panel, with MAF plotted in log scale. Rarest variants are highlighted in b) for comparison. Color coding in a) and b) is the same.

We recapitulate previous results (e.g. [5]) that panel size had a notable effect on imputation accuracy. For a given MAC, *r*^2^ for variants imputed with the largest panel (HRC) is lower than that for variants imputed with the full 1KG panel by ≈ 0.10, the later is in turn lower than that for variants imputed with the European ancestry individuals from 1KG by ≈ 0.04 (difference relatively stable across MAC). However, when conditioning on MAF, the order of panels flips, as the same MAC reflects much smaller MAF in larger panels. Here, *r*^2^ for variants imputed with HRC is higher than that for variants imputed with the full 1KG panel by ≈ 0.05, the later is in turn higher than that for variants imputed with the European ancestry individuals from 1KG by ≈ 0.10.

Sequencing depth of the reference panels on the other hand had only a very small effect. The use of deep coverage 1KG panels increased imputation accuracy typically by < 0.01, compared to the low coverage panel. Note that markers called only in the deep coverage panel are not included in this comparison.

Comparing these empirical result with the theoretical upper bound from our model (figure 4), we observe that the empirical *r*^2^ is much lower than what is predicted under the assumption that only model misspecification affects imputation accuracy. For doubletons imputed with the whole 1KG, the average *r*^2^ is 0.32; for doubletons imputed with the European subset (504 individuals) the average *r*^2^ is 0.41, while the theoretical upper bound is above 0.82 for both sample sizes. While empirical imputation accuracy of rare variants decreases with decreasing MAC across all reference panels, it decreases more rapidly for MAC < 10. This more rapid decrease is mirrored by the rapid decrease of the theoretical upper bound for MAC < 10 observed in the theoretical prediction.

### 3.4 Length between template switches

We now consider another potential source of error in current imputation model: the switch between templates. When we model the target haplotype as a mosaic of templates from the reference, switching between templates can be interpreted as a historical recombination event that breaks the genealogical bond between the target and its current template, i.e., the branches connecting the two leaves in a coalescent tree. The two haplotypes will become practically independent beyond the recombination break point.

Conditional on a known local genealogy, the length to the next recombination break point follows an exponential distribution, with the rate proportional to twice the TMRCA between the target and the template. In practice, the genealogy is unknown, and the length to the next recombination break point is a combination of the distribution of the conditional length and the time to the TMRCA (detailed in Method). Here we compare this mixture distribution with the exponential distribution that is typically assumed in current imputation methods [6, 26].

Comparing the mixture distribution to an exponential distribution with the same mean shows that with genealogy and population history aware modeling, the distribution of length between switches is very similar for small reference samples (*n* = 500) (Figure 6). For larger reference samples (*n* = 20, 000) the mixture distribution has larger variance, with higher density in both extremely short and long intervals but lower density for intermediate ones. If the excess of short no-recombination intervals is not well captured due to model misspecification, we expect to have more suboptimal templates thus higher imputation error rate. When the reference size increases, switches are less frequent as it is more likely to find a template sharing a very recent MRCA with the target.

**Figure 6:**
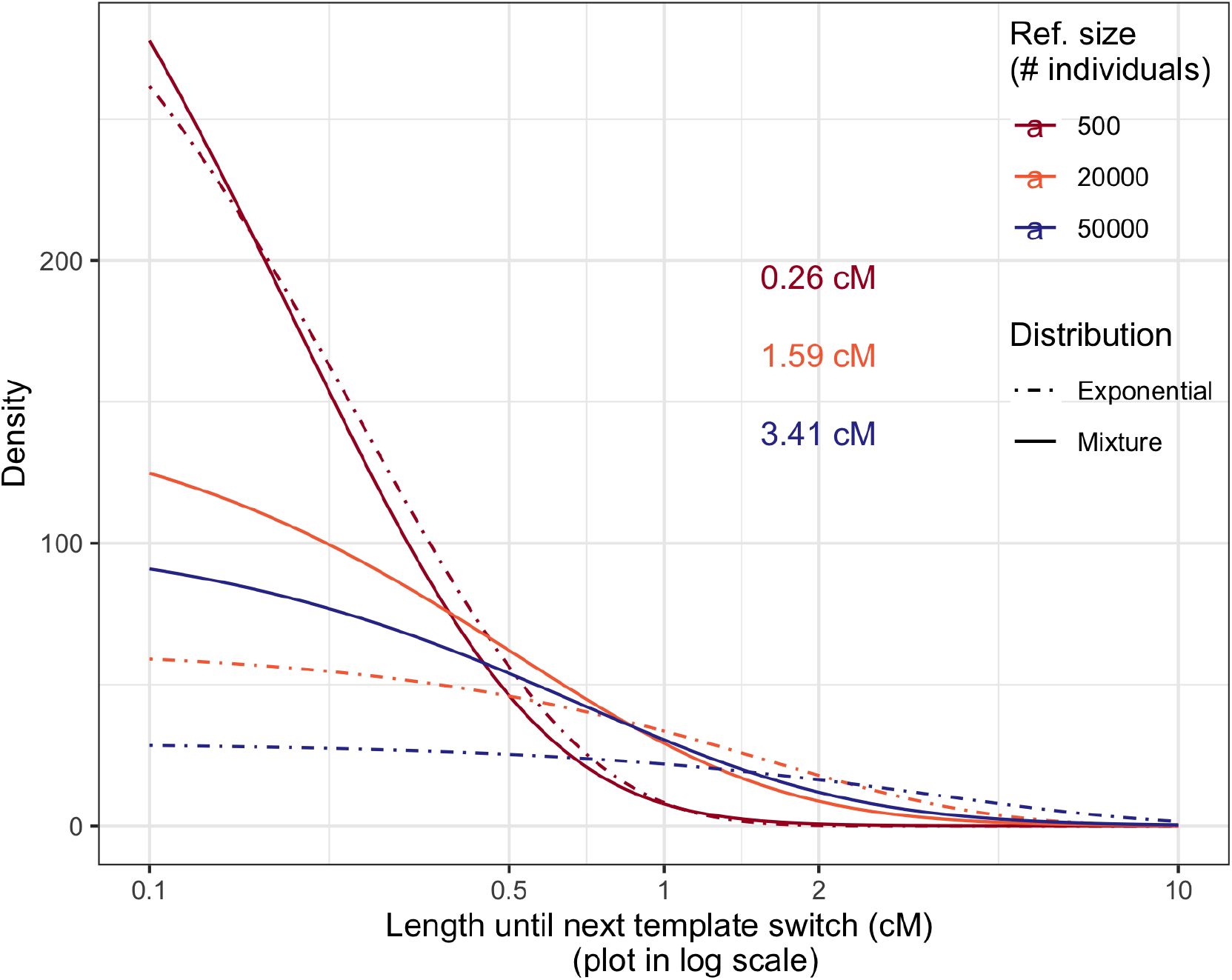
Distribution of the distance before the next recombination event breaking the relation between a pair of haplotypes. The x-axis shows the length in genetic distance (cM). Solid line is predicted by our coalescent model, the dashed line is an exponential distribution with the same mean as we predicted. Colors represent different reference sizes (n) in the number of individuals. The average lengths with reference being n=500, 20000, 50000 are 0.26, 1.59 and 3.41 cM respectively.

## 4 Discussion

The empirical performance of imputation methods has been extensively studied [4, 5, 27, 28], while the theoretical behavior and limit of the underlying framework has not been well characterized, especially for rare variants. We formulate this problem of imputation accuracy into a coalescent model, considering imputing the missing genotypes on one target haplotype by copying from a set of reference haplotypes that share the most recent common ancestor (MRCA) with the target. Our approach identifies two model misspecifications in modern imputation algorithms and explores their impact on our ability to impute rare variants.

First, most imputation algorithms model a single best template as the hidden variable to infer. We show that in 1/3 of the cases, multiple haplotypes are equally good templates for imputing one target in any given reference panel, independent of reference sample size or population history. The resulting model misspecification leads to imputation error when genetic variants are shared by some of the optimal templates but not with the target. We develop analytical expressions for the imputation error resulting from this model misspecification as a function of the derived allele frequency and reference size.

For this purpose, we assume that the imputation algorithm correctly identifies the most closely related haplotypes so that this model misspecification is the only source of error. In this idealized scenario, we observe that for variants observed five or less times in the reference panel, > 8% of variants with non-zero dosage are non-carriers; up to minor allele count 10, the mean *r*^2^ between the imputed dosage and true genotypes < 0.98. Conditional on the derived allele count, this effect is broadly independent of reference panel size: although the probability of falsely imputing carriers of the rare allele decreases as the reference panel size becomes large, the number of true carriers also decreases and the proportion of false carriers over true carriers stays about the same. Including other sources of error that occur in practice further decreases imputation accuracy. Thus the expected *r*^2^ we present here designate the upper bound of achievable mean imputation accuracy for rare variants in current imputation framework.

Such imputation error substantially reduces the power for detecting novel risk variants in an association study based on imputed genotypes. Two scenarios can be considered here: First, single-marker tests of imputed rare variants can be powerful if the case-control data set is much larger than the imputation panel [4]. In this scenario, inaccurately inferred genotypes at a disease-related locus attenuate the allele frequency difference between cases and controls, and *r*^2^ is directly related to this loss of information that compromises the statistical power in association tests. Imputation error rate of 2% ~ 6% leads to 10% ~ 60% increase in required sample size in a single marker test [24]. As a second scenario, imputed rare variants can be aggregated into a single test statistic [2]. In this study design, poorly imputed variants will have an attenuated signal, potentially diluting the signal from better imputed variants. This loss of power can be limited by focusing on imputed variants where misspecification will not impede imputation accuracy, e.g. variants that occur more than five times in the reference.

We consider a second model misspecification: the use of exponential distributions to model the length of contiguous haplotypes without template switches. The true distribution of this length is driven both by the recombination rate and the relationship between the template haplotype and the target haplotype. For smaller reference such as the 1000 Genomes [18] (2504 genomes) this model misspecification has a negligible effect, but for reference samples of the scale of TopMed (53,831 genomes), gnomAD [29] (15,708 genomes), or the Haplotye Reference Consortium [19] (38,821 genomes), the length distribution of shared segments between the target and a single template has a much heavier tails than modelled. This misspecification penalizes both extremely short and extremely long switching intervals, decreasing the probability of finding the optimal templates. This in turn reduces imputation accuracy, especially for low frequency variants beyond the effect of the first model misspecification described above.

The theoretical results we present here are broadly robust to assumptions about population size history in a range reasonable for major human populations. We assume that the imputed region is evolutionary neutral; for loci under selection the impact of the described model misspecifications would likely depend both on the selection model and on the population history. Further, we assume that the reference is from the same homogeneous population as the target sample. If we instead modeled a diverse reference sample, results would depend on the frequency distribution of the imputed variant among the reference samples. For rare alleles, which are typically private to a single population, only members of that single population among the reference would affect imputation accuracy. Our results can be extended to including migration, admixture or selection, as the mathematical derivation and simulation scheme in this work are general for coalescence at a single locus.

In our empirical evaluation of imputation quality using two commonly used reference panel, the accuracy of rare variants is well below theoretical upper bound, suggesting that we have not reached the limit of the Li and Stephen’s framework. Conditional on MAC, imputation becomes less accurate with increasing reference size, suggesting that failure to identify the genealogically best templates is likely the biggest source of error. Identifying these best templates may not always be possible with available marker data, whose resolution is limited by the observed polymorphic sites and local variation of recombination rate. Other factors that are likely to reduce imputation accuracy include error in statistical phasing in the reference or the target haplotypes and departures from the infinite sites assumption (parallel mutations or back mutations). However, as our model predicts, for all reference panels, empirical imputation accuracy decreases much faster for MAC < 10, suggesting that for these low counts the model misspecification modeled here further reduces the ability to correctly impute rare variants.

Our results suggest that potential improvement of the imputation framework may lie in more detailed modeling of the underlying genealogy, especially for extremely rare variants where only a small subset of the reference contributes information about the imputed genotype. For example, suppose the ancestral recombination graph (ARG) including both the target and the reference haplotypes is constructed, the scenarios resulting in fractional dosage could be avoided (up to uncertainty in ARG). However, alleles absent from the reference would still be missed, and there would be uncertainty when all and only the closest reference haplotypes carry an allele: we can only probabilistically decide whether the mutation event or the coalescence event with the target haplotype happens first.

Overall, we identify that the model misspecifications in imputation algorithms limit our ability for imputing rare variants. Such inherent errors reduce the power of single variant tests as well as aggregation tests in studies that impute genotypes in large cohorts, especially if they focus on alleles that are observed only a few times in the reference panel. Beyond improving our understanding of the performance of imputation algorithms, these results point to potential new imputation strategies that help identify new risk variants.

## Acknowledgements

We thank Noah Rosenberg and Jonathan Terhorst for thoughtful comments, and two anonymous reviewers for helpful feedback. Funding for this research was provided by US National Institutes of Health (NIH) US National Institutes of Health grant R01 HG005855.

The authors acknowledge the Michigan Genomics Initiative participants, Precision Health at the University of Michigan, the University of Michigan Medical School Central Biorepository, and the University of Michigan Advanced Genomics Core for providing data and specimen storage, management, processing, and distribution services, and the Center for Statistical Genetics in the Department of Biostatistics at the School of Public Health for genotype data curation, imputation, and management in support of the research reported in this publication.

The following cell lines/DNA samples were obtained from the NIGMS Human Genetic Cell Repository at the Coriell Institute for Medical Research: NA06984, NA06985, NA06986, NA06989, NA06994, NA07000, NA07037, NA07048, NA07051, NA07056, NA07347, NA07357, NA10847, NA10851, NA11829, NA11830, NA11831, NA11832, NA11840, NA11843, NA11881, NA11892, NA11893, NA11894, NA11918, NA11919, NA11920, NA11930. NA11931, NA11932, NA11933, NA11992, NA11994, NA11995, NA12003, NA12004, NA12005, NA12006, NA12043, NA12044, NA12045, NA12046, NA12058, NA12144, NA12154, NA12155, NA12156, NA12234, NA12249, NA12272, NA12273, NA12275, NA12282, NA12283, NA12286, NA12287, NA12340, NA12341, NA12342, NA12347, NA12348, NA12383, NA12399, NA12400, NA12413,, NA12414, NA12489, NA12546, NA12716, NA12717, NA12718, NA12748, NA12749, NA12750, NA12751, NA12760, NA12761, NA12762, NA12763, NA12775, NA12776, NA12777, NA12778, NA12812, NA12813, NA12814, NA12815, NA12827, NA12828, NA12829, NA12830, NA12842, NA12843, NA12872, NA12873, NA12874, NA12878, NA12889, NA12890.NA12890]. These data were generated at the New York Genome Center with funds provided by NHGRI Grant 3UM1HG008901-03S1 (Michael Zody, personal communication).

## A Probability distribution of fractional dosages

In the Method section, we derived the probability of having even the best possible evidence suggesting a wrong allele type. Here we will generalize it to the cases where the most closely related templates have different allele types, resulting in a fractional estimated dosage (figure 2 d).

Let *P*(*ĝ*= *j*/*u, g* = 0|*j*; *n*) be the probability of having *u* equally good optimal templates, *j* of which carrying the mutation, for a target actually carrying the ancestral allele; conditional on observing the derived allele count as *j*. Let (*k′, d′*) denote the numbers of ancestral lines within the size *u* subtree when the branch carrying the mutation starts and ends; (*k″, d″*) denote the numbers of lines in the whole tree at the corresponding time point.

We will outline the calculation of the joint probability (the numerator), while the denominator is the same unfolded frequency spectrum as before. We proceed by the following steps, each subsequent probability is conditional on the previous one, in parallel with that for the special case in the Method.

1. The target haplotype does not coalesce in the first (*n* − *d*) events: *P*_0_(*n* + 1, *d* + 1).
2. A branch of size *u* arises in the reference at the (*n* − *k*)-th event, then remains alone till the (*n* − *d*)-th event to coalesce with that external branch: 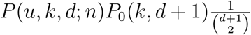.
3. The subtree of size *u* contains a branch of size *j*: *P*(*j*|*u*; *n*) = ∑_(*k′,d′*)_ *P*(*j, k′, d′*|*u*; *n*). (This step only concerns the topology within the subtree)
4. The size-j branch encounters a mutation:*P*(*m* ≥ 1|*j, k′, d′, u, k, n*) = ∑_(*k″,d″*)_ *P*(*m* ≥ 1|*k″, d″*; *n* + 1)*P*(*k″, d″*|*k′, d′, u, k, n*). (Here we need to put the subtree back to the whole size-(*n* + 1) tree to get the branch length and introduce mutation)

Step 1-2) are similar to the three steps in the previous section except for not involving mutation events, we combine them to *P*(*ext, u, k, d*; *n*): an external branch first coalesces at the (*n* − *d*)-th event with an internal branch of size *u* which starts at the (*n* − *k*)-th event. In step 3), *P*(*j, k′, d′*|*u*; *n*) is similar to *P*(*u, k, d*; *n*) in step 2), as *P*(*j, k′, d′*|*u*; *n*) = *P*(*j, k′, d′*; *u*): the probability for a branch of size *j* to start at the (*u* − *k′*)-th event and end at the (*u* − *d′*)-th event.

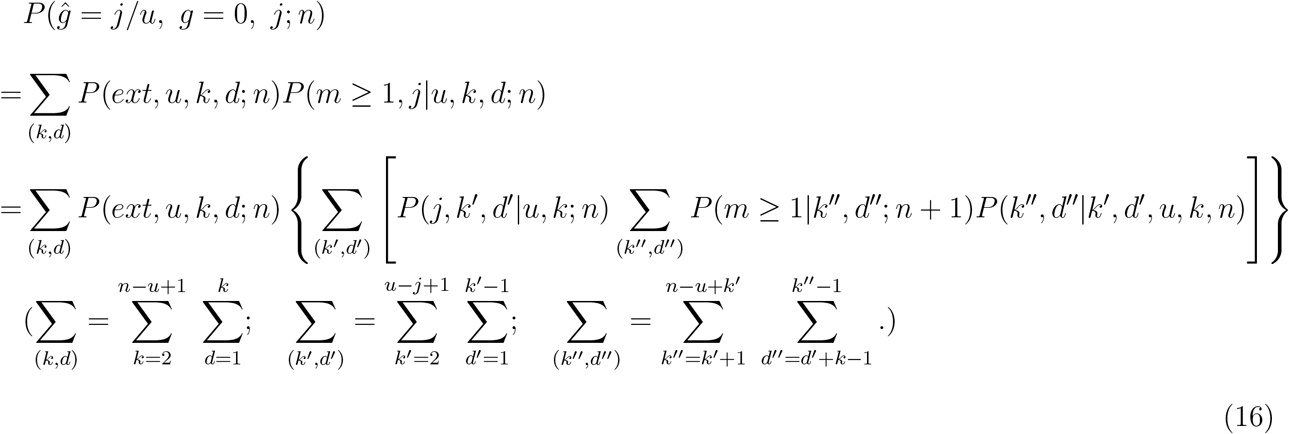

The only component left is *P*(*k″, d″*|*k′, d′, u, k, n*), the link between the topology within the subtree and the branch lengths, which is relative to the whole sample and involves the population size history model. Since coalescent time intervals are measured for the whole tree, (*k″, d″*) tells us when the branch of size *j* starts and ends, which leads to the probability for a mutation to occur.

We divide the coalescent process among the whole reference from *n* lines to *k* lines into three parts: *n* → *k″* → *d″* → *k*. There are (*n* − *k″* − 1)+(*k″* − *d″* − 1)+(*d″* − *k* − 1) unfixed events (since we are conditioning on the three time points, (*k′, k″*), (*d′, d″*) and (*k*)). Within each part, we consider the number of events that has to happen inside the subtree: (*u* − *k′* − 1), (*k′* − *d′* − 1), (*d′* − 1 − 1). With the number of lines fixed, *P*(*k″, d″*|*k′, d′, u, k, n*) is calculated by considering the possible ways to arrange those events in the three time intervals.

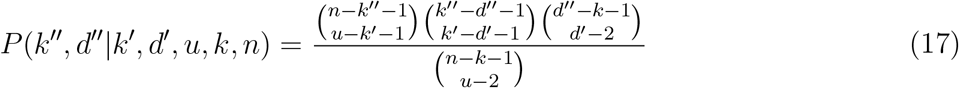

The denominator in (10) comes from removing the constrains introduced by the mutation event. Conditional on (*u, k*), there are in total (*n* − *k* − 1) unfixed events in the whole tree, including (*u* − 2) in the subtree. Since (*k″, d″*) defines the coalescent time in the whole size-(*n*+1) tree, we can calculate the length of the branch carrying mutation by (6): 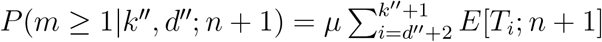.

## B Comparison of opulation growth models

**Figure 7:**
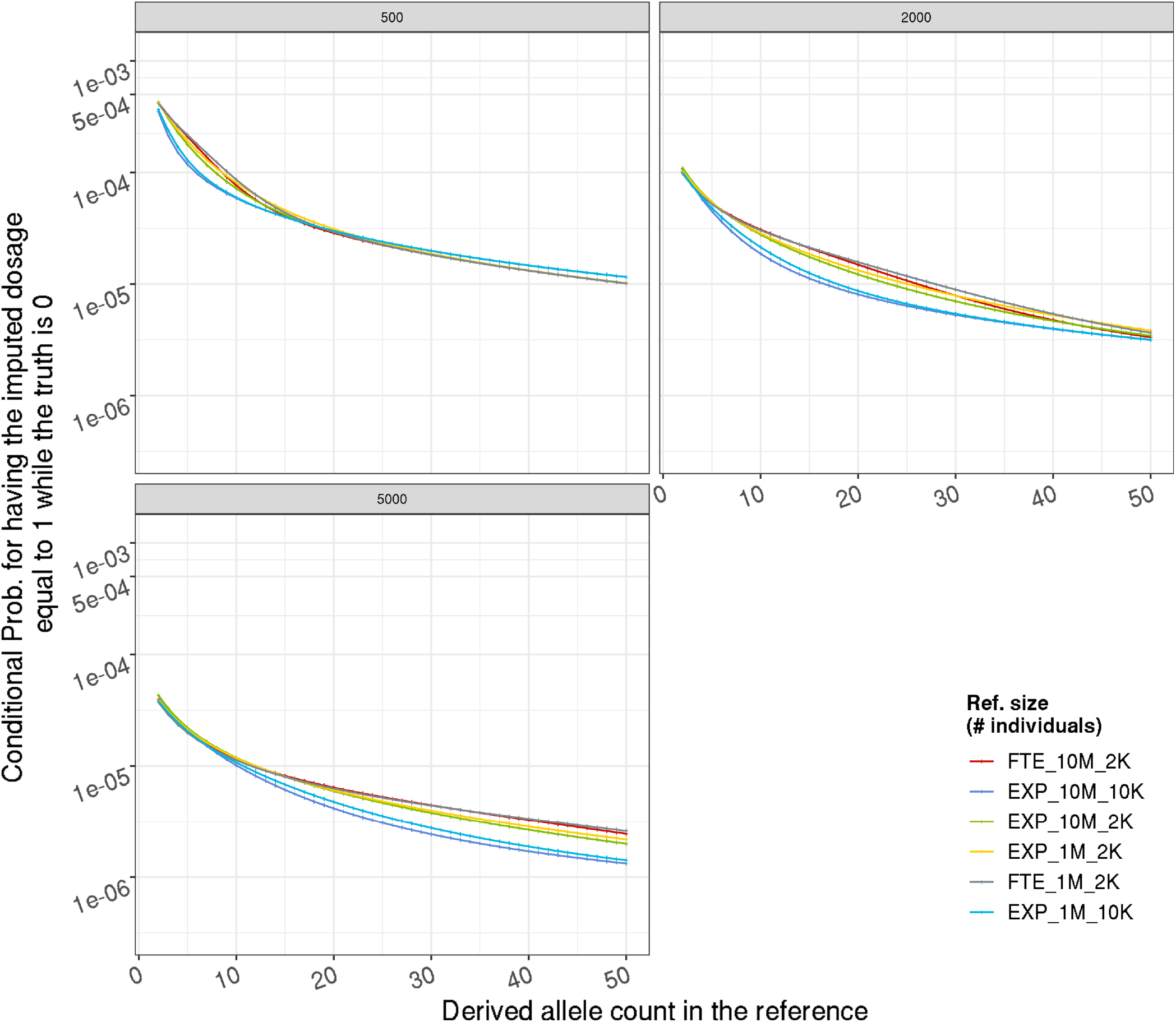
Effect of population growth model on false discovery rate among non-carriers, from analytical calculation. Given a target haplotype carrying the ancestral allele, the probability (y-axis, in log scale) of having all the most closely related reference haplotype in the reference panel carrying the derived allele thus an estimated dosage 1, on the derived allele frequency in the reference (x-axis). Each sub-figure represents one reference size (in individual); each color represents a population growth model. FTE(EXP) 10M 2K: faster-than-exponential (exponential) growth with current day effective population size 10 millions, and a bottleneck with effective population size 2 thousands.

**Figure 8:**
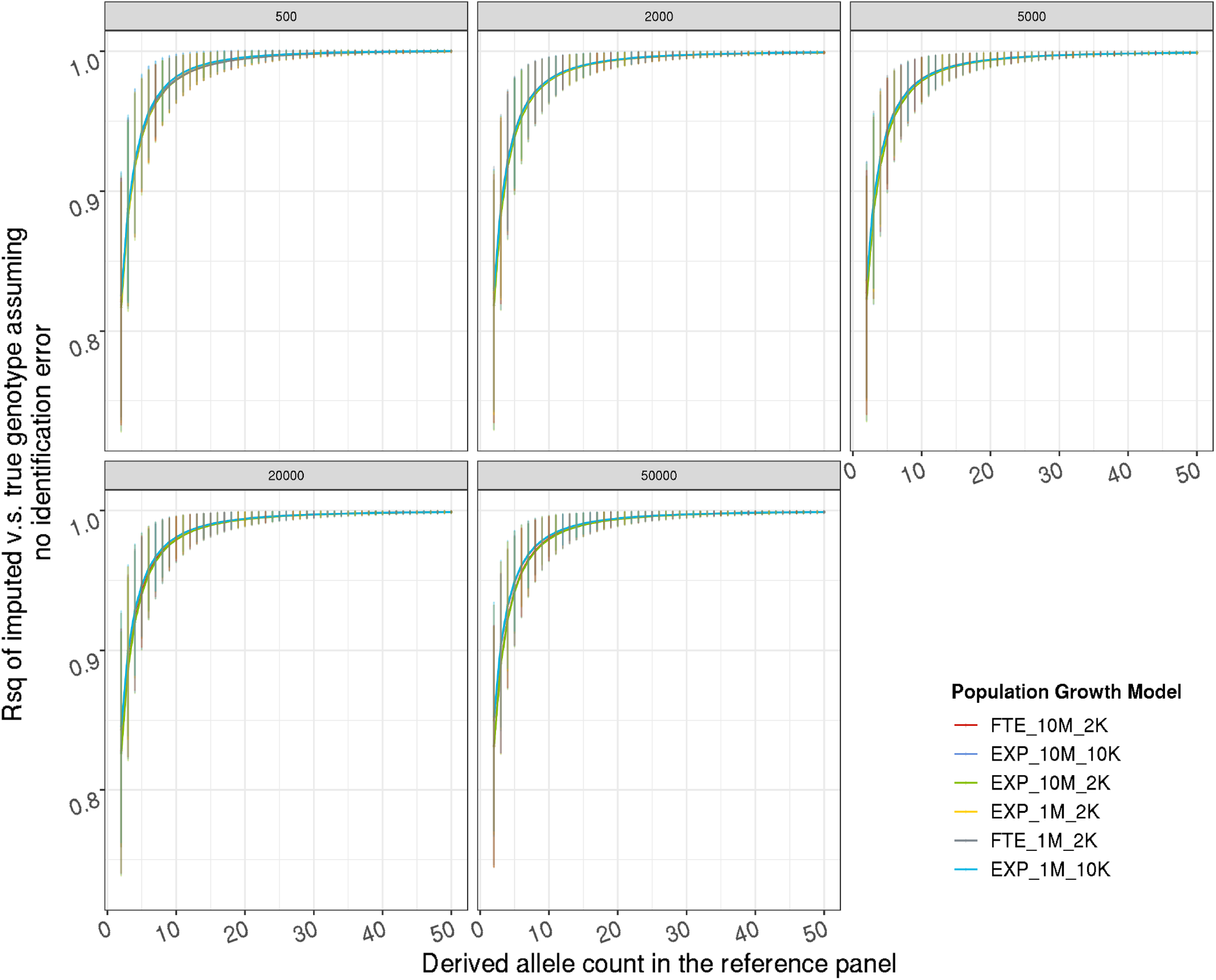
Effect of population growth model on imputation accuracy upper bound (*r*^2^), from coalescent simulation. Y-axis is the average *r*^2^ between imputed dosages and true genotypes without identification error; error bars represent one standard deviation. Each sub-figure represents one reference size (in individual); each color represents a population growth model. FTE(EXP) 10M 2K: faster-than-exponential (exponential) growth with current day effective population size 10 millions, and a bottleneck with effective population size 2 thousands.

## C Genotyping and Sample Quality Control

DNA samples from the blood of Michigan Genomics Initiative (MGI) participants were processed on one of two production batches of a customized Illumina Infinium CoreExome-24 bead array. Genotype calls were produced with the Illumina GenomeStudio 2.0 software operating the Genotyping Module v2.0.4 and the Gentrain clustering algorithm v3.0. Variant and sample level quality control (QC) is detailed as following.

Reads mapping and genotype calling were performed according to the Genome Reference Consortium Human Build 37 (GRCh37), 502,255 variants passed variant level QC. Variants were excluded if (1) bead array probe sequences did not perfectly and uniquely map to the reference (BLAT v.351) [1], (2) variant-level call-rate was below 99%, (3) variant fail Hardy-Weinberg equilibrium exact test with *p* < 10^−^6 in unrelated European samples (PLINK v1.90) [2], (4) GenomeStudio GenTrain score < 0.15 or Cluster Separation score < 0.3, or (4) allele frequency differed between bead array production batches (*p* < 10^−^3, Fisher’s exact test). The genetic ancestry of MGI participants was inferred by projecting MGI samples onto the space created by the first two principal components (PCs) of 938 unrelated samples of the Human Genome Diversity Project (HGDP) reference panel [3] (PLINK). MGI samples were inferred to belong to a HGDP reference population if they fell within a circle drawn around that population in a plot of the PCs. We also lift it over to GRCh38, where 501,607 variants remained.

Only consent individuals were included, and 56,984 samples passed sample level QC. Samples were excluded in QC if (1) genotype-inferred sex is abnormal or did not match the self-reported gender of the participant or the self-reported gender was missing, (2) sample shared a kinship coefficient > 0.45 with another sample with a different identification tag (KING v2.1.3) [4], (3) sample-level call-rate was below 99% or any chromosome had a call-rate bellow 1/5 of the average, or (4) estimated contamination level exceeded 2.5% (VICES) [5].

## Notes

### Competing Interest Statement

The authors have declared no competing interest.

### Summary of Updates

We added a new section on comparing empirical imputation accuracy with the theoretical bound. We added introductory Figure 1 and 2. The manuscript is substantially revised and to appear in Genetics.

